# Genome-wide single-cell perturbation screens with VIPerturb-seq

**DOI:** 10.64898/2026.02.12.705613

**Authors:** Alexandra Bradu, John D. Blair, Erin M. Cumming, Isabella N. Grabski, Isabella Mascio, Junsuk Lee, Cecilia McCormick, Radhika Rathi, Benan Nalbant, Chao Dong, Caleb A. Lareau, Rahul Satija

## Abstract

CRISPR-based screening combined with single-cell sequencing (i.e., Perturb-seq) enables systematic mapping of genetic perturbations to molecular phenotypes. While Perturb-seq is well-suited to profile targeted subsets of regulators, scaling to genome-wide screens presents substantial cost and throughput challenges. Here we introduce VIPerturb-seq, a platform to facilitate routine genome-wide Perturb-seq experiments using probe-based detection workflows. We describe a split probe strategy for detection of genome-wide CRISPR libraries in fixed cells that enables (i) support for phenotypic enrichment of Very Important Perturbations (VIPs) prior to single-cell profiling, and (ii) compatibility with combinatorial indexing workflows to further improve Perturb-seq throughput by 50-fold. Using a genome-wide CRISPRi library (GuEST-List), we demonstrate VIPerturb-seq on three genome-wide screens representing both unbiased and phenotypically enriched workflows. Our results demonstrate how the sensitivity, scalability, and efficiency of VIPerturb-seq can enable both individual labs with targeted research questions and large data generation platforms aiming to construct virtual cells.

## INTRODUCTION

The combination of CRISPR-based pooled screening ^1^ and single-cell profiling has created a powerful framework for causal dissection of cellular regulation. By pairing CRISPR guide RNA (gRNA) capture with single-cell sequencing, technologies like Perturb-seq ^2–4^ can reveal the consequence of many distinct perturbations on transcriptional state in a single experiment. As the multiplexed experimental design removes batch effects across perturbations and controls, these datasets represent a unique, high-resolution, and causal resource to map gene function ^5^ and construct ‘virtual cell’ models ^6^.

However, the power of Perturb-seq remains constrained by significant practical limitations: the number of cells that must be sequenced—and therefore the total experimental expense—scales linearly with the number of perturbations tested. This leads to a fundamental tradeoff where researchers can either preselect a small subset of perturbations to test ^7,8^, losing the ability to identify or characterize novel regulators, or they can attempt unbiased genomewide screens that require profiling millions of cells ^5,9,10^. Unbiased approaches incur substantial cost, much of which is inefficiently allocated towards sequencing perturbations that have little or no phenotypic effect. To fully enable the potential of Perturb-seq for discovery, new approaches are needed that efficiently achieve genome-wide breadth.

Here we introduce VIPerturb-seq to facilitate routine genome-wide single-cell CRISPR screens (Figure 1A). VIPerturb-seq addresses current limitations in two key ways. First, it is compatible with fixed samples and therefore supports diverse modes of phenotypic enrichment, allowing researchers to enrich for cells exhibiting a phenotype of interest (Very Important Perturbations) before single-cell profiling. Therefore, the cost-limiting library preparation and sequencing steps are disproportionately focused on cells with desired phenotypes, dramatically reducing the number of cells to profile without needing to pre-specify the perturbations of interest. Second, the workflow incorporates combinatorial indexing where transcripts are dually labeled with both a well and droplet barcode, facilitating the generation of massively scalable datasets.

**Figure 1.**
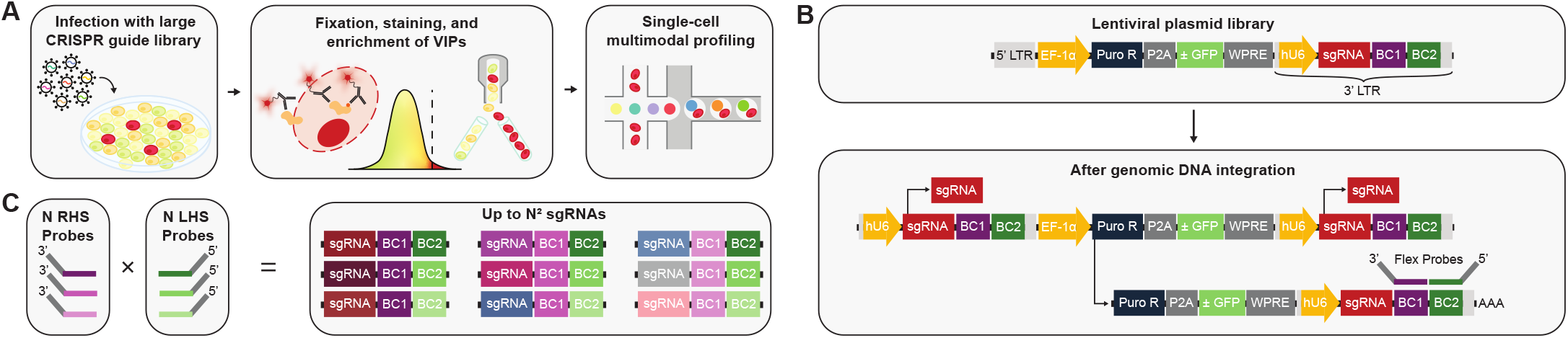
Overview of VIPerturb-seq. **A)** Schematic of the VIPerturb-seq experimental workflow. Cells are transduced with a large pooled CRISPR guide library, optionally subjected to phenotypic enrichment (Very Important Perturbations; VIPs) using fixation and intracellular antibody staining, and subsequently profiled using multimodal single-cell sequencing. **B)** Schematic of the lentiviral construct design. Each CRISPR sgRNA is paired with a synthetic barcode transcript compatible with probe-based detection. **C)** Each barcode sequence is defined by a unique combination of left-hand side (LHS) and right-hand side (RHS) barcode halves, which are independently recognized by split probes.

Aiming to enable genome-wide pooled CRISPR screens within a fixed-cell single-cell RNA-seq workflow, VIPerturb-seq is built around probe-based detection of known transcripts, for example with the 10x Genomics Flex platform. Targeted RNA detection improves data quality and enables compatibility with fixed samples ^11^, but is not natively suited for capturing tens of thousands of synthetic sequences such as a genome-wide gRNA library. To overcome this challenge, VIPerturb-seq associates each perturbation with a dual-barcode system designed for split-probe detection, enabling unambiguous assignment of more than 50,000 gRNA identities using fewer than 500 custom probes.

We show that this framework supports fixation and intra-cellular antibody staining, phenotypic enrichment of perturbed cells, and downstream single-cell profiling — allowing new discoveries from a genome-scale perturbation screen after profiling just 12,000 single cells. Moreover, even in the absence of phenotypic enrichment, we use combinatorial barcoding with Flex v2 (Flex Apex) to profile 440,000 cells per single 10x reaction lane. This workflow enables a 50-fold improvement in throughput, allowing the molecular consequences of perturbing every human gene to be profiled in a routine experiment, and is accompanied by a 65% increase in detected genes per cell compared to conventional Perturb-seq approaches. Together, these features make VIPerturb-seq a versatile and scalable solution for routine genome-wide single-cell perturbation screening.

## RESULTS

### Overview of VIPerturb-seq workflow

We sought to develop VIPerturb-seq as a platform for genome-wide Perturb-seq screens that is fully compatible with diverse strategies for phenotypic enrichment. Enrichment for molecular phenotypes, particularly intracellular proteins or RNA markers, typically requires cellular fixation and permeabilization, processes that substantially degrade RNA and limit compatibility with conventional reverse-transcription (RT)-based single-cell workflows. We therefore needed an alternative to RT-based RNA capture and turned to probe-based single-cell profiling, which is specifically designed for fixed samples with degraded RNA. For example, the Flex technology from 10x Genomics uses a split-probe architecture, in which each target locus is recognized by two independent probes, the left-hand side (LHS) and right-hand side (RHS) probes, that must both hybridize and ligate in order for the molecule to be detected. This split-probe design increases specificity and makes Flex compatible with diverse upstream processing workflows, including fixation and permeabilization, followed by intracellular antibody or RNA FISH probe staining ^12^. However, because Flex relies on targeted probes, directly detecting tens of thousands of unique gRNA protospacer sequences would require constructing an enormous probe panel that would scale linearly with library size, and would need to be independently repeated for different types of genetic perturbations (i.e., CRISPRi vs. CRISPRa libraries). We therefore pursued an alternative and generalizable perturbation readout strategy that maintains compatibility with custom probe-based capture while supporting genome-wide screens.

To achieve this, we linked each perturbation’s identity with a synthetic barcode sequence (Figure 1B), allowing Flex probes to detect the barcode rather than the sgRNA protospacer itself. Previous studies have highlighted that barcode–guide mispairing can occur due to recombination events during lentiviral packaging, particularly when the genomic distance between the barcode and protospacer is large ^13^. To minimize recombination between these elements, we used a CROP-seq-based vector ^14^ expressing a Pol III-driven sgRNA and a Pol II-transcribed barcode (Figure 1B), and we reduced the genomic distance between protospacer and barcode to less than 100 bp (compared to 2,500 bp in previous approaches ^13^). As in previous work where synthetic barcode sequences were read out via imaging ^15^, we also introduced three CR backbone variants (Supplementary Figure 1A), with each gRNA randomly utilizing one of these three backbones, to further limit homology-mediated rearrangements ^2^.

The use of a synthetic barcode allows sequences to be flexibly designed to optimize capture and scalability. We therefore implemented a split-barcode system in which each perturbation is defined by a unique combination of two barcode halves. This allows us to leverage the split-probe chemistry of the Flex assay to independently recognize the two halves by custom LHS and RHS probes (Figure 1C). This design substantially reduces the requirements for custom probe synthesis compared to direct sgRNA capture, as a library of *n* perturbations can be directly read out using a total of 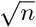 probe pairs. As these sequences are not tied to CRISPR-specific architecture, identical barcode and probe sets can be reused across distinct sgRNA libraries. They may even be paired with entirely different synthetic elements, such as ORF overexpression constructs ^16^, variant libraries ^17^, or clonal lineage barcodes ^18^, making this a generalizable strategy for detecting user-introduced sequences.

### Robust barcode recovery and gRNA assignment

We first performed a pilot experiment to assess whether VIPerturb-seq could induce perturbations, capture guide identities with high sensitivity, and generate high-quality multimodal single-cell data. We designed a library of 100 sgRNAs targeting known and putative mTOR regulators, along with non-targeting controls, for which we had previously generated Perturb-seq data ^19^. We cloned a library of constructs where each gRNA was associated with a unique LHS/RHS combination of one of 10 LHS probes and one of 10 RHS probes to facilitate detection with our custom probe set. To test compatibility across multiple biological systems, we transduced a pool of K562, HEK293, and HAP1 cells with lentivirus encoding this library, and performed a VIPerturb-seq experiment using the 10x Flex v1 workflow (Figure 2A). To assess our capability to generate multimodal data, we introduced barcoded antibodies targeting phosphorylated RPS6 (pRPS6) using our previously introduced FlexPlex workflow ^19^, allowing us to quantify intracellular protein levels of this mTOR signaling marker by sequencing antibody-derived tags (ADT).

**Figure 2.**
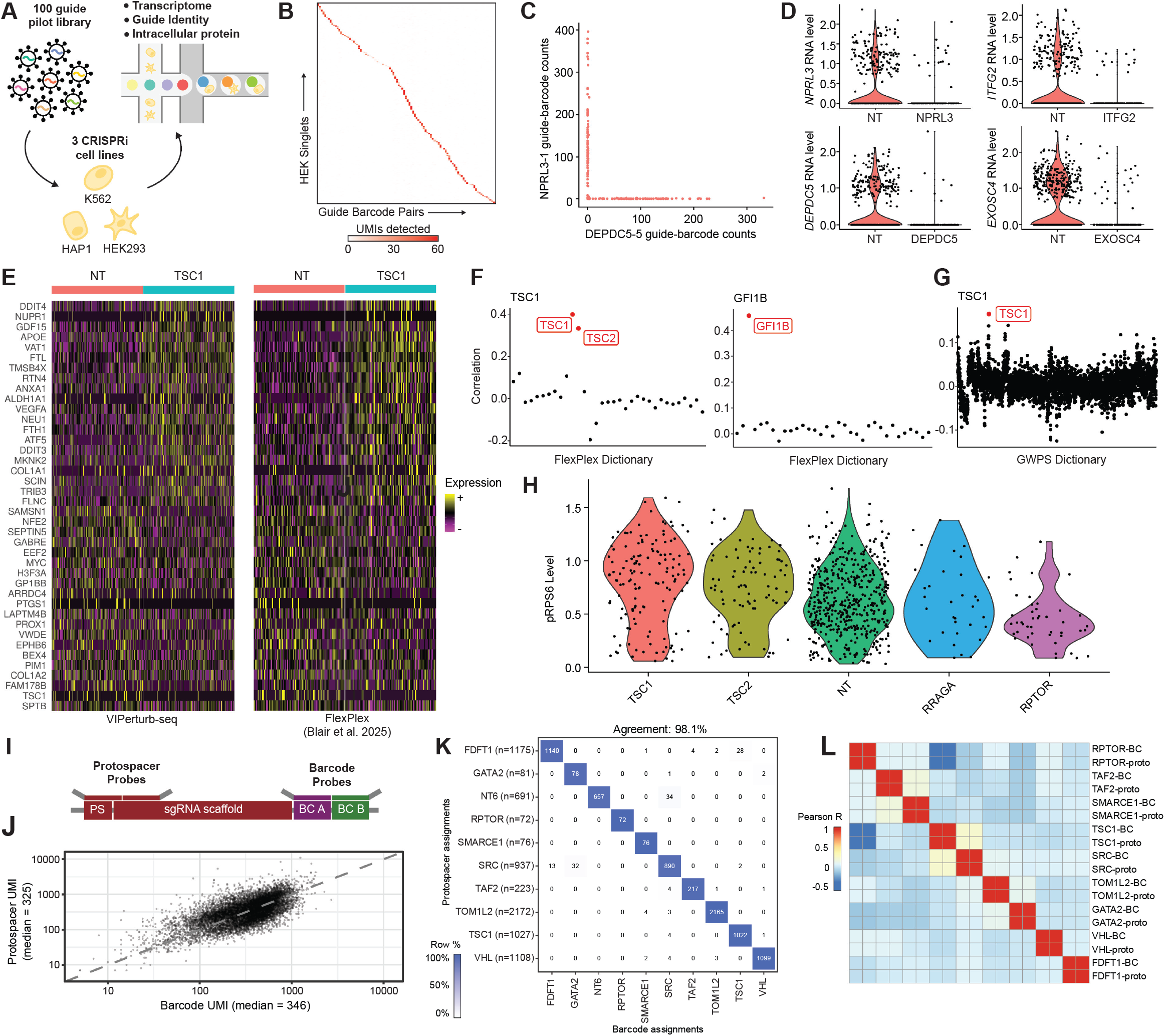
Pilot VIPerturb-seq experiments to assess gRNA barcode recovery and multimodal profiling. **A)** Schematic illustrating pilot experimental design, performed across three cell lines to assess generalizability. **B)** Heatmap of UMI counts of each guide-barcode in all HEK293 cells called as singlets. Cells are ordered by assigned guide identity. **C)** Barnyard plot comparing counts of two distinct guide-barcode pairs per cell. **D)** Violin plots of target RNA levels for four representative target genes. **E)** Heatmap of upand downregulated genes between TSC1-perturbed and non-targeting K562 cells, in both this dataset (left) and an independent FlexPlex dataset. **F)** Long Island City plots for TSC1-perturbed (left) and GFI1B-perturbed (right) K562 cells, plotting mapping similarity using RNA fingerprints learned from the FlexPlex dataset depicted in (E). Identified matches are colored in red. **G)** Long Island City plot for TSC1-perturbed K562 cells using RNA fingerprints learned from GWPS. Identified matches are colored in red. **H)** Violin plots of pRPS6 antibody-derived tag levels across perturbations of mTOR pathway members in K562 cells. **I)** Schematic illustrating the recombination benchmarking experiment, including protospacerand barcode-targeting probes for guide detection. **J)** Scatter plot comparing barcode and protospacer UMI counts in cells called as gRNA singlets by both modalities. **K)** Confusion matrix comparing protospacer-based and barcode-based guide assignments across 7,562 cells called as singlets in both modalities. **L)** Heatmap of Pearson correlations between perturbation RNA fingerprints learned independently using protospacer-based and barcode-based guide assignments.

Across all three cell types, we observed strong recovery of barcode sequences and robust assignment of gRNA identities (Figure 2B,C). In HEK293, we observed a median of 52 UMI for the most abundantly detected sgRNA within an individual cell, followed by a median of 1 UMI for the second-most abundant sgRNA (Supplementary Figure 2A-C). This enabled confident assignment of perturbations to more than 95% of cells (with 87% classified as singlets). Comparable performance metrics were observed in K562 and HAP1 cells (Supplementary Figure 2A-C). As expected for robust gRNA recovery and accurate assignment in CRISPRi screens, gRNA levels were mutually exclusive and associated with transcriptomic knockdown of the assigned target gene (Figure 2C-D).

Consistent with previously observed high mRNA sensitivity from the Flex assay ^11^, we obtained 6,700 RNA UMI/cell even at low sequencing saturation (10,300 reads/cell). To confirm the accuracy of mRNA and gRNA data, we compared the results from differential expression between *TSC1* versus non-targeting (NT) assigned cells in our dataset with results from an independent 10x Flex experiment in which guides were read out through direct gRNA protospacer capture ^19^. The transcriptional signatures showed high concordance across the two datasets (Figure 2E). We further benchmarked perturbation signatures by applying RNA fingerprinting ^20^, which aims to match the phenotypic effects of perturbed cells to a reference Perturb-seq dictionary. VIPerturb-seq cells showed specific and robust fingerprinting to the correct target perturbation either when using our previous Flex Perturb-seq dictionary (Figure 2F), or alternately, the Genome-Wide Perturb-seq (GWPS) dataset (Figure 2G) generated in K562 cells ^5^. The resulting matches confirm that VIPerturb-seq accurately recapitulates expected gene-expression consequences of perturbation. Perturbation of mTOR regulators also resulted in expected changes to measured intracellular pRPS6 levels, a downstream marker of mTOR signaling, confirming the ability to simultaneously measure intracellular ADTs (Figure 2H).

### Direct benchmarking demonstrates highly concordant protospacer and guide-barcode assignments

Accurate perturbation assignment requires each synthetic barcode to remain linked to its intended sgRNA throughout pooled lentiviral production and transduction. To directly measure barcode-guide mispairing, we performed a dedicated benchmarking experiment using nine targeting sgRNAs and one non-targeting control. Following transduction of K562 CRISPRi cells, we used custom Flex probes to detect both the sgRNA protospacer and its associated synthetic barcode in the same cells (Figure 2I). This gave us the opportunity to benchmark both the capture sensitivity and identity of the recovered transcripts, providing a precise measure of the lentiviral recombination rate.

Both readouts showed comparable and cell-correlated recovery at single-cell resolution (Figure 2J). The sensitivity for barcode vs. protospacer recovery (346 vs. 325 median UMI/cell) is notable because protospacer probes capture both the highly expressed Pol III functional sgRNA and the barcoded Pol II transcript, whereas barcode probes capture only the Pol II transcript ^14^. Therefore, these results reflect the ability for VIPerturb-seq probes to achieve enhanced hybridization and capture of optimized synthetic barcode sequences.

We next compared protospacer-based and barcode-based guide identity calls. Across 7,562 cells called as singlets in both modalities (Supplementary Methods), assignments agreed in 98.1% of cases, corresponding to a guide–barcode discordance rate of 1.9% (Figure 2K). To determine whether the rare discordant assignments affected biological interpretation, we independently learned transcriptomic fingerprints for each perturbation using either protospacer-based or barcode-based assignments. Fingerprints for matched perturbations were nearly identical, with a median Pearson correlation of 0.97 (Figure 2L), and differentially expressed genes between perturbed and nontargeting cells exhibited identical patterns (Supplementary Figure 2D). Thus, barcode-based assignment not only accurately recovers sgRNA identity, but also yields the same inferred transcriptomic consequences of perturbation as direct protospacer capture. Together, these results support the effectiveness of our strategies to minimize recombination during lentiviral production and transduction, and establish VIPerturb-seq synthetic barcodes as an accurate and sensitive readout of sgRNA identity.

### GuEST-List: a genome-wide barcode library

Having established the technical feasibility of VIPerturb-seq, we next sought to extend the method to true genomewide scale. We therefore designed a genome-wide CRISPRi perturbation library by adapting the Dolcetto library A, which contains up to three sgRNAs targeting each annotated human protein-coding gene, along with non-targeting controls ^21^. Instead of restricting profiling to a subset of genes, the genome-wide Dolcetto library is agnostic to cell type or transcriptional state (Figure 3A). This breadth comes at the cost of scale: with 57,050 sgRNAs, per-perturbation cell requirements necessitate sequencing very large numbers of cells.

**Figure 3.**
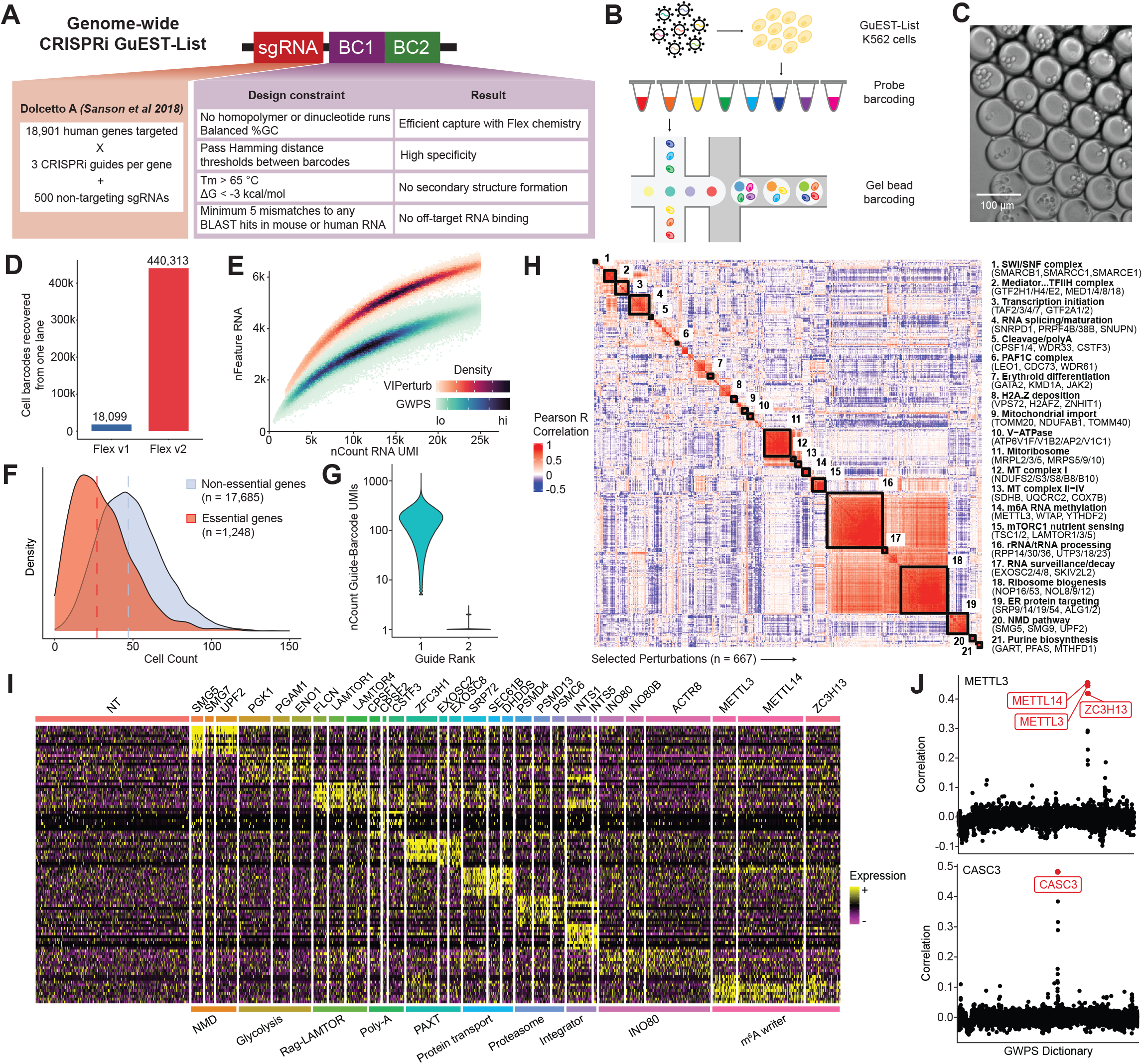
Genome-wide VIPerturb-seq screening with GuEST-List. **A)** Summary of computational parameters governing barcode sequence design for GuEST-List. **B)** Schematic illustrating combinatorial barcoding approach to improve cellular throughput with 10x Genomics Flex v2 chemistry. **C)** Image of multicellular GEMs formed using the Flex v2 workflow, where the combination of well and droplet barcodes enable single-cell demultiplexing. **D)** Number of cell barcodes recovered per 10x lane using the Flex v1 and v2 workflows. **E)** Density scatterplot showing detected UMIs versus genes per cell in VIPerturb-seq and GWPS ^5^. **F)** Density plot of cell coverage across perturbations (dashed line indicates median). **G)** Violin plot of guide-barcode UMIs per cell for the top versus second-most abundant guide. **H)** Correlation matrix and clustering of filtered perturbation fingerprints, with manual annotation of 21 functionally enriched clusters and representative genes. **I)** Heatmap of top upregulated genes for ten groups of biologically related perturbations. **J)** Long Island City plots for METTL3 and CASC3-perturbed cells. Matches to GWPS are colored in red.

To enable single-cell readout of this library using VIPerturb-seq, we generated a corresponding set of synthetic barcode sequences and imposed a series of stringent sequence-level constraints to ensure robustness at genome-wide scale (Figure 3A). First, all barcode halves were required to maintain a minimum Hamming distance from one another, allowing single-base mismatches arising from sequencing error, probe misligation, or transcriptional noise to be distinguished from true barcode identities. Second, to promote uniform hybridization and ligation efficiency across the panel, candidate sequences were constrained to a narrow range of GC content and melting temperature. We further filtered sequences using *in silico* structural analyses to eliminate those predicted to form hairpins, self-dimers, or other secondary structures that could interfere with probe binding ^22^. Third, to prevent off-target hybridization, we excluded any sequences with appreciable similarity to the human or mouse transcriptomes, ensuring that barcode probes bind exclusively to synthetic transcripts rather than endogenous RNA. Finally, to minimize unintended cross-pairing within the split-probe architecture, the probe-binding regions corresponding to the LHS and RHS halves were designed to have minimal complementarity across the full probe set. The resulting optimized barcode space represents 238 LHS and 240 RHS sequences, encoding 57,120 possible barcode sequences. We provide these (Supplementary Tables) as a reusable and standardized resource for detecting large synthetic libraries using probe-based single-cell workflows.

We refer to this genome-wide perturbation library as GuEST-List (Guides for Enrichment-based Screening of Targets Library), reflecting its broad inclusion of known protein-coding genes, even though only a subset of perturbations may emerge as VIPs in any given screen. Below, we demonstrate how the GuEST-List library enables either unbiased profiling of all perturbations (facilitated by scalable profiling with Flex v2), or alternatively, focused profiling of VIPs following phenotypic enrichment.

### Pre-indexing facilitates massively scalable profiling

While GuEST-List enables the detection of genome-wide gRNA libraries, naïvely applying standard single-cell sequencing workflows remains logistically and financially challenging. For example, perturbing 20,000 genes at a depth of 500 cells per gene would require 10 million cells. With conventional 10x scRNA-seq, the costs for library preparation and sequencing for this experiment surpass $1,000,000. We therefore next focused on reducing per-cell costs by increasing cellular throughput without compromising data quality.

To address library preparation costs, we adapted VIPerturb-seq for compatibility with the 10x Genomics Flex v2 (Apex) chemistry, which introduces a pre-indexing step prior to microfluidic droplet generation (Figure 3B). In this workflow, cells are first split across multiple wells, hybridized with transcriptome and VIPerturb-seq barcode probes, and labeled with a well-specific oligonucleotide barcode. During subsequent droplet encapsulation, multiple cells can be jointly loaded and barcoded together, while transcripts remain uniquely assignable to individual cells based on their well and droplet barcodes (Figure 3B,C). Increasing the number of pre-indexing wells therefore directly increases effective cellular throughput in a single library preparation reaction (“lane”).

As a proof of concept for unbiased genome-scale screening, we performed a VIPerturb-seq experiment using the GuEST-List library in K562 cells. Cells were pre-indexed across 24 wells and processed across two Flex v2 lanes, targeting approximately 420,000 cells per lane. This represents a greater than 20-fold expected increase in throughput relative to conventional droplet-based single-cell workflows. The Flex v2 platform provides 384 pre-indexing reactions, supporting up to and beyond 1 million cells per lane, demonstrating that increases of 50-fold or greater in throughput are achievable with the same workflow.

Sequencing was performed using an Ultima Genomics UG100 instrument, and we recovered approximately 880,000 cell barcodes across two lanes (Figure 3D). Robustness of gRNA assignment exceeded our pilot experiment, with a median of 160 UMIs per cell for the most abundant gRNA, and 1 UMI/cell for the second-most abundant (Figure 3G). We were able to assign a perturbation identity to 97% of cells, with 81% classified as gRNA singlets. As an additional successful control, we did not observe evidence of reads mapping to the small fraction (70 of 57,120) of possible GuEST-List LHS/RHS combinations that were unused in our library design.

Transcriptomic data quality was similarly high, with a median of 15,100 RNA UMIs and 5,300 detected genes per cell. Despite a sequencing depth that is far from saturation (25,000 reads/cell), both metrics compared favorably to the previously published genome-scale Perturb-seq ^5^ (Figure 3E), reflecting a median increase of 30% UMI/cell and 65% genes/cell and demonstrating that the improvements in scalability for VIPerturb-seq are also accompanied by improvements in sensitivity.

We note that this genome-wide dataset represents a proof-of-concept experiment with limited cellular depth, given that these cells are distributed across 57,050 sgRNAs (median of 14 cells/gRNA, 42 cells/targeted gene) (Figure 3F). As expected, we also observed a further depletion of perturbations previously annotated as essential (Figure 3F). Despite this, we found that the data were nevertheless sufficient to support multiple orthogonal analyses that together demonstrate successful genome-scale perturbation profiling. First, we applied our previously developed RNA fingerprinting framework to estimate the transcriptional effect of each perturbation, even when per-gene differential expression is underpowered ^20^. Using this framework, we identified evidence for target gene knockdown in 88% of perturbations (Supplementary Methods). We also identified 6,724 perturbations whose inferred fingerprints passed QC filtering, indicating broader transcriptomic effects beyond depletion of the target gene.

Second, we examined the structure of these inferred fingerprints by clustering perturbations based on the similarity of their transcriptomic fingerprints (Supplementary Methods). As expected from high-quality Perturb-seq data, this analysis revealed strong and biologically coherent clustering of perturbations with known related functions (Figure 3H). For example, we observed clear grouping of perturbations associated with a wide variety of distinct regulatory processes including nonsense-mediated decay, mTOR nutrient sensing, polyadenylation, the RNA exosome, and m^6^A methylation (Figure 3H). Shared perturbation responses were observed both by unsupervised clustering of RNA fingerprints (Figure 3H), and by visualizing single-cell expression levels for perturbations that are known to be functionally related (Figure 3I).

Third, we assessed the correspondence between perturbation fingerprints learned in this experiment and reference fingerprints previously computed from a gold-standard genome-wide Perturb-seq dataset (GWPS) ^5^. Successful matching between datasets indicates that perturbations induce shared transcriptional responses despite differences in experimental platform and sequencing chemistry. In some cases, matching occurred at the level of protein complexes or pathways, such as METTL3 matching to a cluster of m^6^A writer components (Figure 3J). In other cases, we observed precise one-to-one correspondence, such as CASC3 matching specifically to CASC3 (Figure 3J). Taken together, these results show that VIPerturb-seq assigns gRNA identities to cells, successfully identifies both target gene knockdown and downstream transcriptomic effects, and learns perturbation signatures that recapitulate existing technologies with improved per-cell data quality and substantially increased throughput.

These results highlight the applicability of VIPerturb-seq to enable genome-wide screening workflows. However, even with dramatically reduced library preparation costs, performing genome-wide experiments still incurs substantial costs for sequencing millions of cells, many of which may exhibit no noticeable phenotype. Reducing library preparation costs therefore represents only a first step to improve scalability, requiring further developments to enable routine and cost-effective genome-wide Perturb-seq.

### Phenotypic enrichment of VIPs

VIPerturb-seq enables a complementary strategy to dramatically reduce the number of profiled cells in a genomewide screen: instead of sequencing all cells, only those exhibiting a phenotype of interest are selectively enriched and profiled. We applied this approach to the cytoskeletal intermediate filament protein vimentin (VIM), which forms a dynamic network regulating cell mechanics ^25^, organization ^26^, and stress responses ^27^ and whose abundance must be tightly controlled to prevent cytoplasmic stiffening and disruption of cell division and signaling ^28^.

To identify negative regulators of VIM at genome-wide scale, we transduced K562 cells with the GuEST-List library, fixed, permeabilized, stained for intracellular VIM, and isolated the top 3% of VIM-high cells by FACS. These enriched cells were then profiled using VIPerturb-seq in a single 10x Flex v1 lane, yielding 12,000 sgRNA singlets (Figure 4A). We note that vimentin is most often studied in the context of the epithelial–mesenchymal transition (EMT), where its upregulation accompanies largescale transcriptional and phenotypic remodeling. By instead performing this screen in K562 cells, which do not undergo EMT, we were able to interrogate regulatory mechanisms controlling VIM abundance that are independent of EMT-associated cell state transitions.

**Figure 4.**
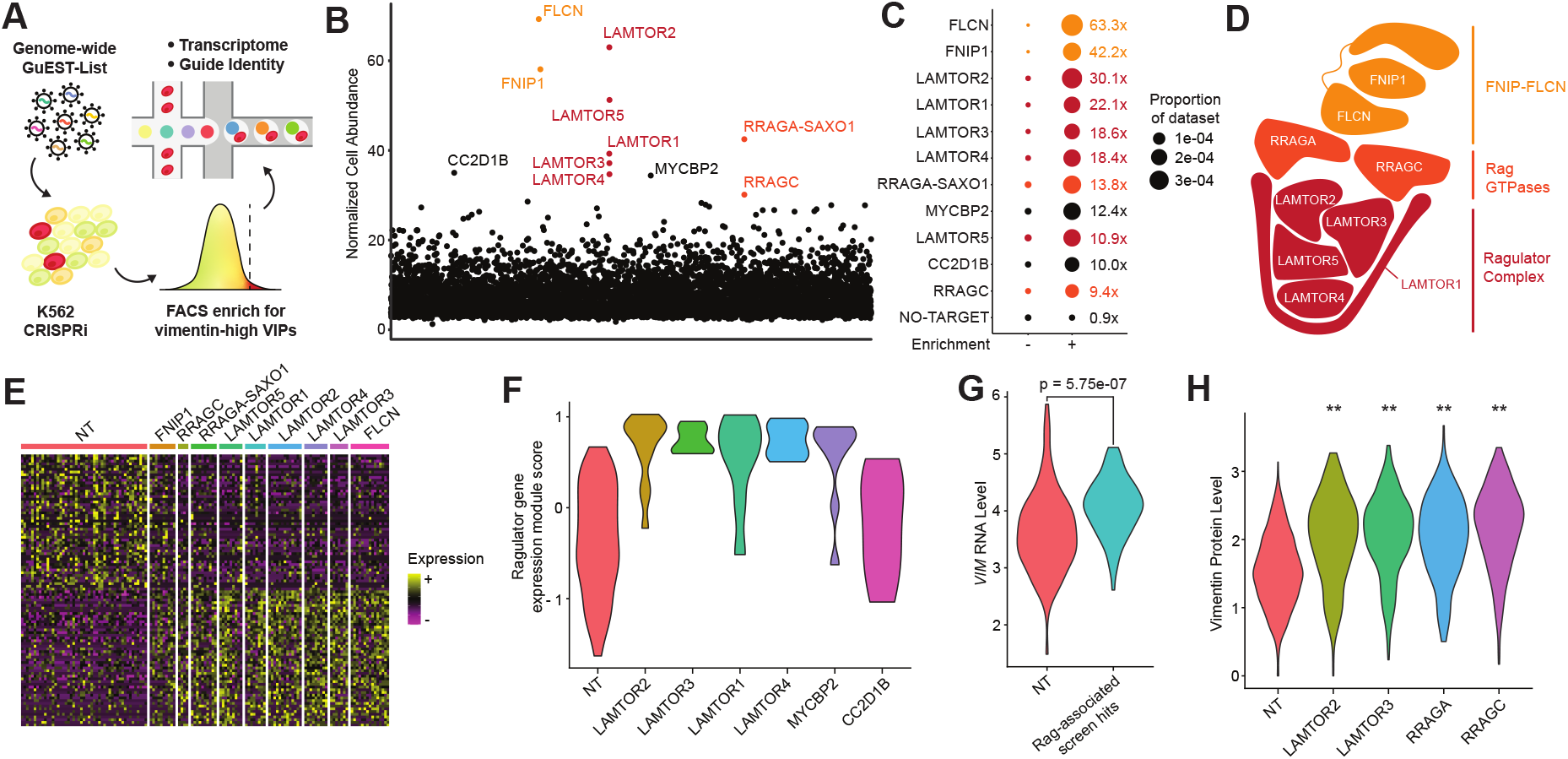
VIPerturb-seq reveals a role for the Rag–Ragulator-FLCN complex in vimentin regulation. **A)** Schematic illustrating the VIPerturb-seq workflow used to screen for negative regulators of vimentin. **B)** Abundance plot illustrating top perturbations recovered from single-cell libraries derived from the vimentin-high, phenotypically enriched FACS fraction. **C)** Fold enrichment of perturbations in the vimentin-high FACS fraction relative to the non-enriched genome-wide screen, with point size indicating the fraction of cells assigned to each perturbation. **D)** Schematic representation of the structural association of Rag–Ragulator-FLCN complex members, adapted from ^23^ and ^24^. **E)** Heatmap of up- and downregulated genes between Rag–Ragulator-FLCN complex-perturbed cells and non-targeting cells. **F)** Violin plots of Ragulator gene module scores across perturbations enriched in vimentin VIPerturb-seq. **G)** Violin plots of *VIM* mRNA expression in non-targeting cells and Rag–Ragulator-FLCN complex-perturbed cells. **H)** Violin plots of vimentin protein levels (normalized intracellular ADT) across perturbations measured in an independent FlexPlex experiment. ^**\*\***^ indicates *P <* 10^*−*27^.

We analyzed the resulting data in two complementary ways. First, we identified candidate regulators solely by considering the sgRNA identities of cells in the enriched fraction, ranking perturbations by their normalized abundance compared to background abundance in the plasmid library (Supplementary Methods). This analysis revealed a highly specific enrichment for components of the Ragulator complex, which governs lysosomal amino acid sensing and activation of mTORC1^29^ (Figure 4B), but to our knowledge has not been previously established to regulate VIM expression. When comparing our enriched to non-enriched screens, we found that phenotypic enrichment increased relative cell recovery for these perturbations by 10 to 63-fold, consistent with the approximately 33-fold enrichment expected in our top 3% sorted bin (Figure 4C).

Strikingly, this analysis recovered the entire Rag– Ragulator–FLCN complex (Figure 4D). Our identified regulators included five canonical Ragulator subunits (LAMTOR1–5), which together form the lysosomal scaffold required for amino acid–dependent mTORC1 recruitment^30^. Both Rag GTPases (RRAGA and RRAGC), which function as a heterodimeric molecular switch to control lysosomal localization of mTORC1 in response to amino acid availability, were strongly enriched. We also identified the FLCN–FNIP1 subcomplex, which functions as a GTPase-activating factor for the Rag GTPases and is required for proper cycling of Rag activity ^31^. These identified regulators represented 9 of the top 11 hits. Moreover, we found that enrichment was highly specific to this nutrient-sensing axis, without enrichment of mTOR itself or other core mTORC1 components.

As VIPerturb-seq provides single-cell transcriptomic profiles for each enriched cell, we next examined the molecular signatures associated with the identified regulators. We first derived a transcriptional signature from perturbations of canonical Rag–Ragulator–FLCN complex members, which exhibited a clear and coherent gene-expression response (Ragulator module) that was conserved across known regulators (Figure 4E). Having established this signature, we then used it to characterize the mechanisms of action of enriched regulators that are not canonical complex members. Perturbation of *MYCBP2*, an E3 ubiquitin ligase previously implicated in mTOR signaling ^32^, induced transcriptional responses that were nearly indistinguishable from those of Ragulator and Rag GTPase perturbations (Figure 4F). Perturbation of CC2D1B, which participates in the mitotic reformation of the nuclear envelope ^33^, did not lead to upregulation of this module, suggesting a potentially alternative regulatory role. Therefore, the transcriptional data accompanying VIPerturb-seq can be used to help understand the mechanism of newly identified regulators, though additional experiments are needed to validate MYCBP2 and CC2D1B, and exclude potential false positive or off-target effects.

As the Rag–Ragulator–FLCN complex represents a putatively novel regulator of vimentin abundance, we validated this result across two modalities. We first examined RNA expression levels of *VIM* in cells perturbed for the identified regulators. While RNA abundance provides an imperfect proxy for regulation of a single protein, pooling cells across perturbations revealed a significant increase in *VIM* expression relative to non-targeting controls (Figure 4G). As a second validation, we examined an independent FlexPlex dataset in which members of the Rag– Ragulator complex were perturbed and ADTs reflecting intracellular VIM protein levels were directly measured ^19^ (Figure 4H). In each case, perturbation of Ragulator complex members was associated with clear upregulation of intracellular VIM, providing reproducible and independent support for the VIPerturb-seq screening result.

### RNA-based phenotypic enrichment identifies regulators of ncRNA processing

Having demonstrated phenotypic enrichment based on intracellular proteins, we next asked whether VIPerturb-seq could isolate phenotypes defined by RNA. Many cellular states lack suitable antibody markers but can be identified by specific transcripts or transcriptional signatures. RNA surveillance provides one example: snoRNA host transcripts such as *SNHG3* and *DANCR* are highly transcribed, yet their processed transcripts are rapidly degraded by the RNA exosome ^34,35^ (Figure 5A). Prior work has highlighted U1 snRNP recognition and the balance between nuclear degradation and export as key determinants of RNA fate ^36–40^. However, how nuclear surveillance selectively targets transcripts for degradation remains an area of active research, and the factors that influence this process remain incompletely understood. Screening cells based on RNA-defined phenotypes should enable the systematic identification of these regulators and provide a deeper understanding of selective ncRNA turnover.

**Figure 5.**
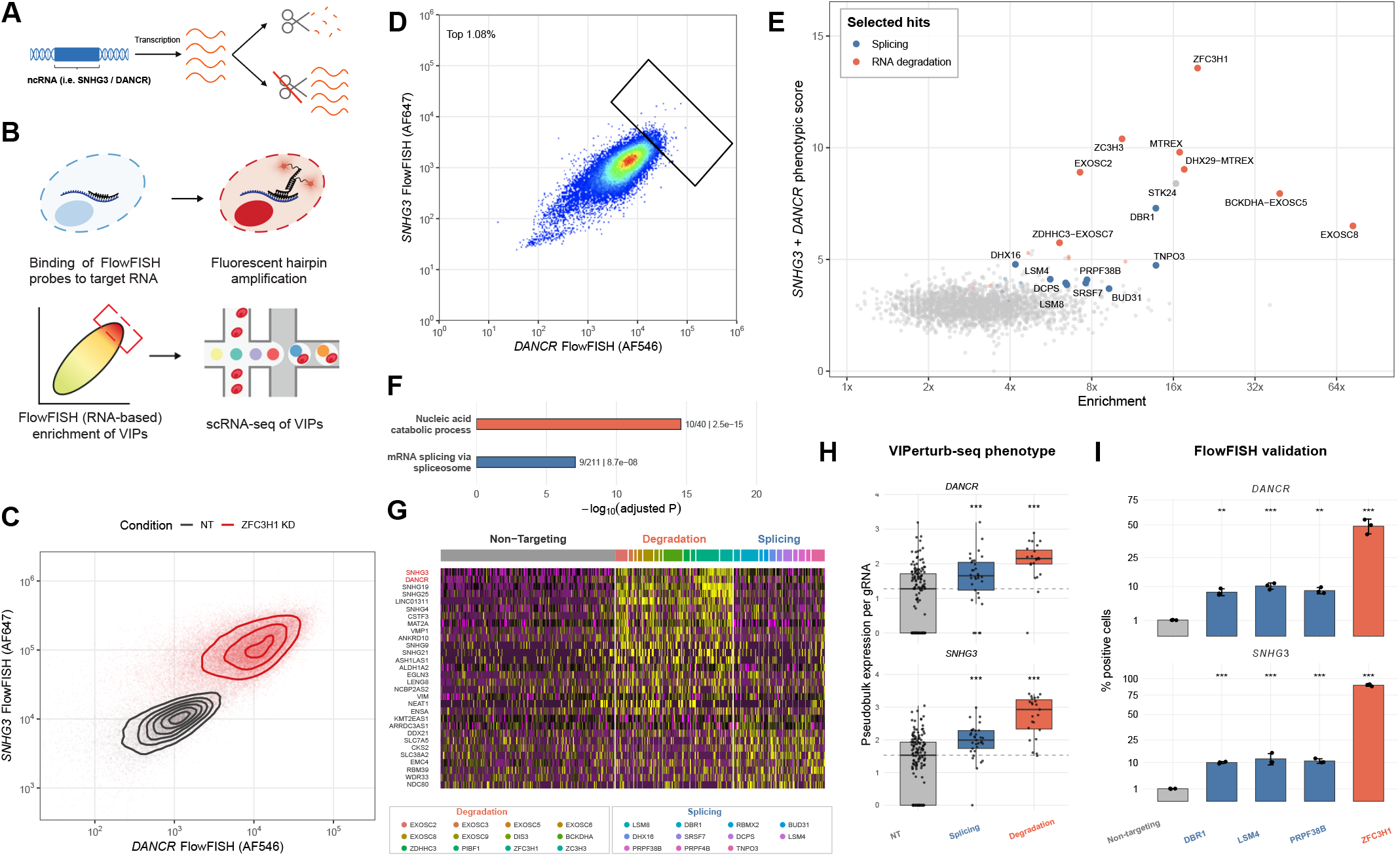
RNA-based phenotypic enrichment identifies regulators of ncRNA abundance. **A)** Schematic of ncRNA accumulation following disruption of nuclear RNA surveillance. **B)** Schematic illustrating the combination of VIPerturb-seq and PERFF-seq for genome-wide screening of regulators of *DANCR* and *SNHG3*. **C)** Flow-cytometry data and density contours of *DANCR* and *SNHG3* in non-targeting and ZFC3H1-knockdown cells. **D)** Gate used to isolate cells with the highest combined *DANCR* and *SNHG3* signal (top 1.08%). **E)** Top screen hits jointly prioritized by enrichment relative to the plasmid library and a molecular phenotype score representing total *DANCR* and *SNHG3* expression across cells carrying each perturbation (Supplementary Methods). **F)** Gene ontology enrichment among the top 50 prioritized perturbations. **G)** Heatmap of selected transcriptional signatures in cells carrying non-targeting guides or RNA-degradation and splicing perturbations. The two ncRNA targets are highlighted in red. **H)** Normalized pseudobulk *DANCR* and *SNHG3* expression by gRNA. Points denote individual gRNAs, boxes summarize distributions, and dashed lines mark non-targeting medians. ^***^ indicates *P <* 0.001 (two-sided Wilcoxon rank-sum test). **I)** Independent FlowFISH validation of selected regulators. Bars show mean percentages of RNA-positive cells, points represent three replicates, and error bars show standard deviations. Y axes use a square-root scale. Stars indicate two-sided contrasts with the non-targeting control from a pooled linear model (Supplementary Methods; ^**^ *P <* 0.01; ^***^ *P <* 0.001).

To accomplish this, we considered recently developed technologies that sort cells based on their expression of RNA markers. For example, PERFF-seq uses HCR-FlowFISH with target-specific probes to convert a cell’s expression of selected RNA transcripts to a fluorescent signal, isolates defined RNA states by FACS, and profiles the sorted cells using 10x Flex ^12,41^. Since both VIPerturb-seq and PERFF-seq utilize 10x Flex for single-cell profiling, we reasoned that the two workflows could be combined (Figure 5B). As an initial test, we performed HCR-FISH using probes targeted to *DANCR* and *SNHG3* in K562 cells followed by FACS. We confirmed that CRISPRi-mediated knockdown of ZFC3H1, a key adapter within the poly(A) tail exosome targeting (PAXT) complex ^34,35^, led to an increase in the expression of both ncRNA transcripts, consistent with sharply reduced degradation (Figure 5C).

To identify other regulators, we transduced K562 cells with GuEST-List and sorted the 1% of cells with the highest combined *SNHG3* and *DANCR* signal as VIPs to identify additional regulators of ncRNA expression (Figure 5D; Supplementary Figure 3A). The sorted population was profiled using 10x Flex, with custom transcriptome probes included to quantify *SNHG3* and *DANCR*, as these ncRNAs are not included in the standard transcriptome-wide panel. We confirmed that the PERFF-seq workflow, which includes additional steps for HCR FlowFISH probe staining and polymer stripping, did not interfere with GuEST-List barcode detection: among 29,379 cells in the top 1% ncRNA-high fraction, 26,956 (91.8%) received at least one gRNA identity and 21,644 (73.7%) were classified as gRNA singlets. These singlets contained a median of 153 total guide-barcode UMIs per cell (Supplementary Figure 3B), and were retained for downstream analysis.

The single-cell profiles allowed candidate regulators to be evaluated using two complementary measurements. For each gene, we calculated sgRNA enrichment in the joint-high population relative to its abundance in the plasmid library and a molecular phenotype score based on the summed *SNHG3* and *DANCR* expression across cells carrying that perturbation. We combined the percentile ranks of these measurements to prioritize perturbations that were both enriched by sorting and associated with the expected molecular phenotype (Figure 5E). As expected, the topranked hits included our positive control ZFC3H1 and multiple components of the PAXT complex and the nuclear exosome. Several additional top-ranked perturbations targeted loci immediately adjacent to known RNA-surveillance genes, including BCKDHA–EXOSC5, DHX29– MTREX, and ZDHHC3–EXOSC7, consistent with local effects of CRISPRi ^42^. STK24 was the only highly ranked hit unrelated to RNA processing, but is likely explained by promiscuous off-target activity recently reported for an STK24-targeting CRISPRi gRNA ^43^.

While recovery of RNA-degradation machinery represents an expected positive control, our primary goal was to identify other processes that may help the surveillance machinery recognize and route transcripts for degradation. We therefore performed gene ontology enrichment on the top 50 jointly prioritized hits. After the expected enrichment for nucleic acid catabolic processes, which included 10 known members, the second strongest nonredundant signal was mRNA splicing via the spliceosome (Figure 5F). In addition to the 9 annotated spliceosome hits in this category, we identified PRPF38B, a Prp38-family protein that organizes protein interactions during spliceosome activation, and TNPO3, the nuclear import receptor for serine/arginine-rich splicing factors ^44,45^. Notably, none of the splicing-associated hits were canonical components of the U1 snRNP; instead, they spanned diverse stages of the broader splicing pathway. Previous work has shown that U1 snRNP can recruit PAXT and LENG8, influencing both direct degradation and nuclear export ^38–40^. Our findings suggest that, for these targets, the relationship between splicing and ncRNA turnover extends beyond U1mediated transcript recognition.

This finding is supported by multiple distinct lines of evidence. Examining the transcriptomic effects of perturbation in our VIPerturb-seq data revealed shared responses to exosome-associated and splicing-associated regulators, with partial overlap among ncRNAs, including *DANCR* and *SNHG3* (Figure 5G). Increased ncRNA expression was supported by both screening readouts: spliceosome-associated hits were enriched across multiple independent sgRNAs (Supplementary Figure 3C) in the sorted ncRNA-high population (Figure 5E) and consistently increased *DANCR* and *SNHG3* expression across multiple independent sgRNAs (Figure 5H). In both cases, the spliceosome-associated hits therefore represented an intermediate tier, exhibiting reproducible effects above non-targeting controls but below the strongest positive control perturbations in both screening enrichment and ncRNA upregulation. This demonstrates that VIPerturb-seq goes beyond recovering the strongest canonical regulators and can identify multiple tiers of regulators across a range of phenotypic strengths in the same experiment.

Finally, we independently validated selected regulators using individual sgRNAs. Motivated by the design of our original screen, we used non-targeting control cells to establish baseline levels of expression for both *DANCR* and *SNHG3* using FlowFISH and set positive gates such that 1% of non-targeting cells were positive for each ncRNA. Perturbation of ZFC3H1, the lariat debranching enzyme DBR1, and the spliceosomal factors LSM4 and PRPF38B increased the proportion of cells positive for both transcripts across three replicate experiments (Figure 5I), with splicing-associated perturbations again showing a reproducible intermediate phenotype. Together, these results demonstrate that VIPerturb-seq can use a multiplexed RNA phenotype to enrich and prioritize perturbations from a genome-wide library. These results connect disruption of mRNA splicing to the accumulation of *DANCR* and *SNHG3*, consistent with a model where productive splicing marks or licenses these transcripts for decay ^40^. More generally, the compatibility of the PERFF-seq and VIPerturb-seq workflows establishes RNA-defined phenotypes as a flexible basis for single-cell genome-wide perturbation screening. As both *DANCR* and *SNHG3* represent multi-exonic long ncRNAs, repeating this approach with different classes of ncRNAs with different structural properties represents an exciting direction for future work.

## DISCUSSION

In this study, we introduce VIPerturb-seq, a platform designed to facilitate routine genome-wide single-cell perturbation screens. VIPerturb-seq associates each perturbation with a two-component synthetic barcode that can be detected by split probe pairs. This design allows genome-wide CRISPR sgRNA libraries to be readily detected, and more broadly, represents a general framework for probe-based detection of large synthetic sequence libraries introduced into single cells regardless of perturbation modality.

While there have been multiple recent efforts to reduce the cost of scRNA-seq and Perturb-seq workflows ^46–50^, a central advantage of VIPerturb-seq is its reliance on probe-based, RT-free RNA detection. Eliminating reverse transcription enables compatibility with phenotypic enrichment strategies that require fixation and permeabilization, such as intracellular protein staining or RNA-based labeling ^12^, while also facilitating massively scalable combinatorial indexing. These advantages are accompanied by a boost in sensitivity in both mRNA and sgRNA detection compared to traditional workflows. The combination of superior data quality, scalability, and cost efficiency— implemented entirely using widely available commercial single-cell infrastructure—positions VIPerturb-seq as a broadly applicable technology for single-cell perturbation screens.

Phenotypic enrichment represents a particularly powerful aspect of this framework. Using intracellular protein enrichment, which would not otherwise be possible with conventional 3’ scRNA-seq, we identified and validated the Rag–Ragulator–FLCN complex as a regulator of vimentin abundance (Figure 4). By combining VIPerturb-seq with HCR-FlowFISH ^12^, we also identified and validated new regulators of an ncRNA-defined phenotype that cannot, by definition, be accessed through protein-based enrichment (Figure 5). In both experiments, phenotypic enrichment enabled discoveries from genome-wide screens after profiling only approximately 20,000 cells—a scale compatible with a routine single-cell experiment. Moreover, we anticipate that the ability to enrich cells based on expression of a single transcript or a composite RNA signature could greatly expand the phenotypic landscape accessible to single-cell perturbation screens. As scRNA-seq continues to identify transient cell states and subtle regulatory programs, flexible RNA cytometry should broadly generalize to enriching these populations.

The modularity of VIPerturb-seq and GuEST-List also offers key practical advantages. Performing Perturb-seq on pre-selected groups of targeted regulators requires project-specific synthesis of gRNA libraries and matched capture probes, which can be both costly and time-consuming. In contrast, the GuEST-List CRISPRi genome-wide library can be reused across biological systems and questions, with phenotypic enrichment determining which perturbations are ultimately interrogated in any given screen. Moreover, alternative perturbation libraries such as CRISPRa, ORF, or variant libraries can be read out using an identical probe panel when paired with the same GuEST-List barcode set. Together, these features reduce experimental overhead, promote standardization across studies, and enable efficient reuse of reagents.

As single-cell perturbation screens continue to emerge as one of the fastest-growing data modalities in biology, improving their accessibility will be essential for maximizing their impact. By combining phenotypic enrichment, combinatorial indexing, and scalable probe-based perturbation readout in a unified platform, VIPerturb-seq provides an attractive solution for routine genome-wide Perturb-seq. We anticipate that this framework will help broaden adoption of single-cell perturbation screening and accelerate the systematic mapping of gene and variant function across biological contexts.

## ACKNOWLEDGEMENTS

We acknowledge Peter Smibert, Paul Lund, Andrew Kohlway, Reuben Saunders, and Will Allen for helpful discussions in the design of VIPerturb-seq. We acknowledge Ultima Genomics and the New York Genome Center Sequencing core for their assistance and sequencing support. We acknowledge Alex Marson, Ronghui Zhu, Paul Datlinger, Orit Rozenblatt-Rosen, Spyros Darmanis, Pratiksha Thakore, Joachim de Jonghe, and Aziz Al’Khafaji for additional helpful discussions. I.N.G. is the Kenneth G. Langone Quantitative Biology Fellow of the Damon Runyon Foundation (DRQ-21-24). This work was supported by the Chan Zuckerberg Initiative (EOSS5-0000000381, HCA-A-1704-01895 to R.S.), the NIH (RM1HG011014, 1OT2OD033760-01, 4UH3NS132139-03 to R.S., R33CA302491 to C.A.L.), and the MacMillan Center for the Study of the Non-Coding Cancer Genome. We acknowledge the use of the Memorial Sloan Kettering Cancer Center Flow Cytometry Core Facility, funded in part through the NIH/NCI Cancer Center Support Grant P30 CA008748.

## DATA AND REAGENT AVAILABILITY

Seurat is made available as an open-source R package at https://github.com/satijalab/seurat. RNA finger-printing is made available as an open-source R package at https://github.com/satijalab/rna-fingerprinting. Seurat objects representing all single-cell experiments generated for this manuscript are available on Zenodo: https://doi.org/10.5281/zenodo.18460279.

The CRISPRi GuEST-List v1 and CRISPRi GuEST-List GFP v1 libraries are available from Addgene under accession numbers #248188 and #248189.

We include sequences for oligonucleotide probes, library inserts and custom primers (Supplementary Tables), and step-by-step protocols for VIPerturb-seq (Supplementary File) as supplementary material, and at https://satijalab.org/viperturb-seq.

---

**Supplementary Figure 1.**
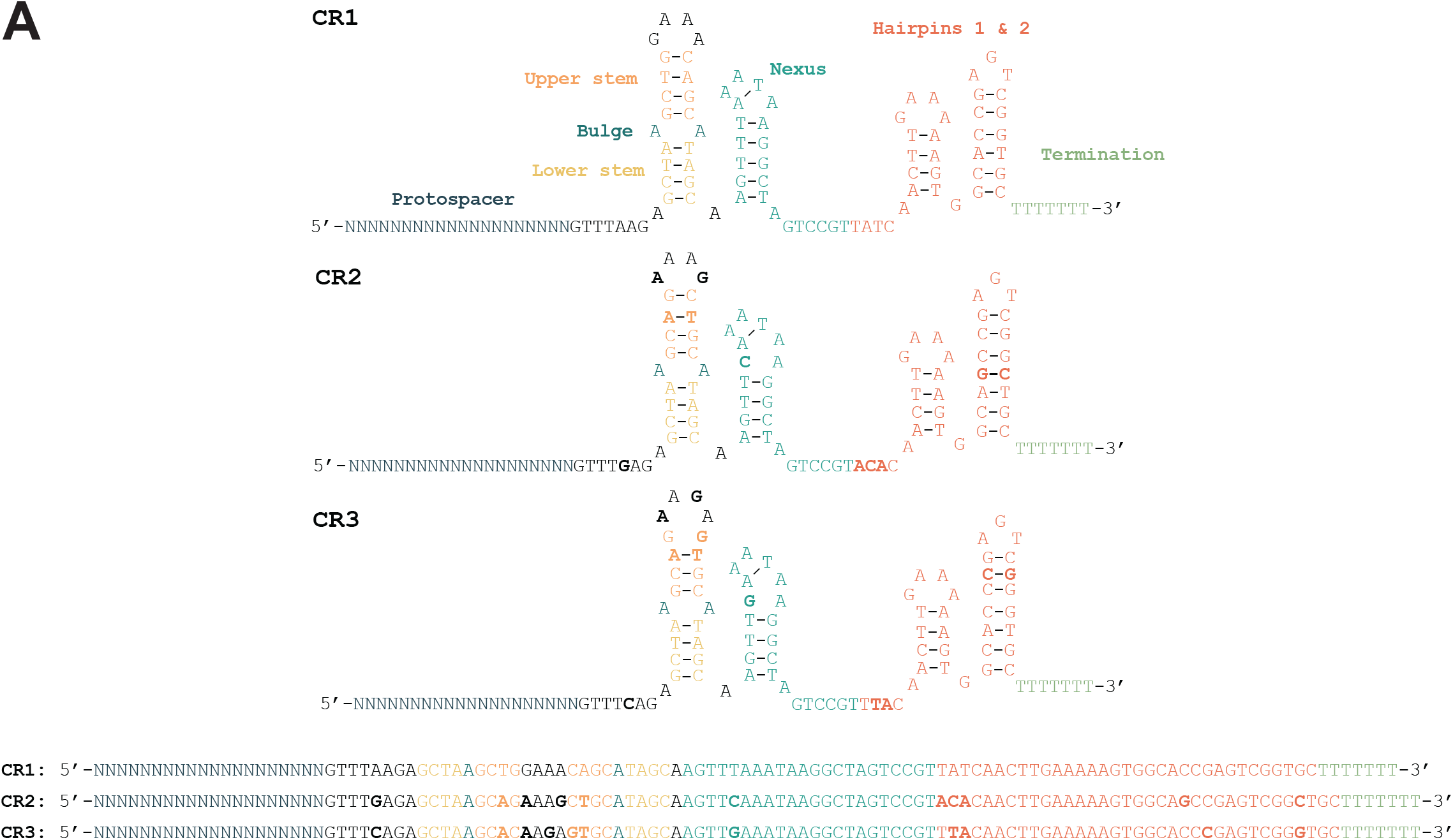
**A**) Schematic representation of three variant sgRNA constant regions used in GuEST-List, as originally described in Adamson et al. 2016 2. While the overall structure and function of the sgRNA is preserved across the three variants, the highlighted changes ensure no continuous stretches of homology between the scaffolds exceeding 20 bp.

**Supplementary Figure 2.**
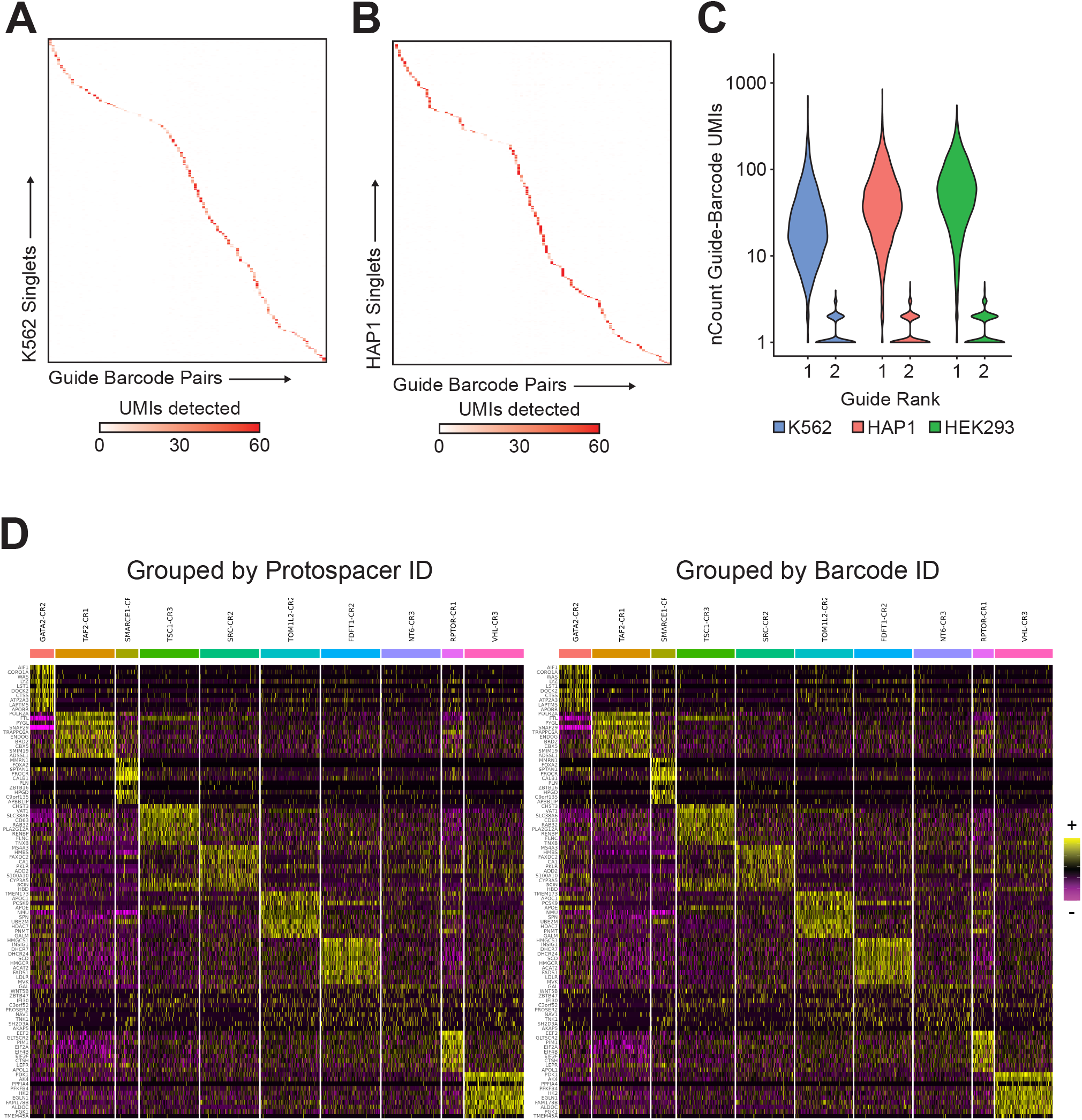
**A)** Heatmap of guide-barcode UMI counts in K562 cells called as singlets. Cells are grouped by assigned guide identity. **B)** Heatmap of guide-barcode UMI counts in HAP1 cells called as singlets. Cells are grouped by assigned guide identity. **C)** Distributions of UMI counts for the highest- and second-highest-abundance guide-barcode in K562, HAP1, and HEK293 cells. **D)** Gene-expression heatmaps for cells called as singlets in both modalities in the recombination benchmarking experiment, grouped using either protospacer-derived or barcode-derived guide assignments.

**Supplementary Figure 3.**
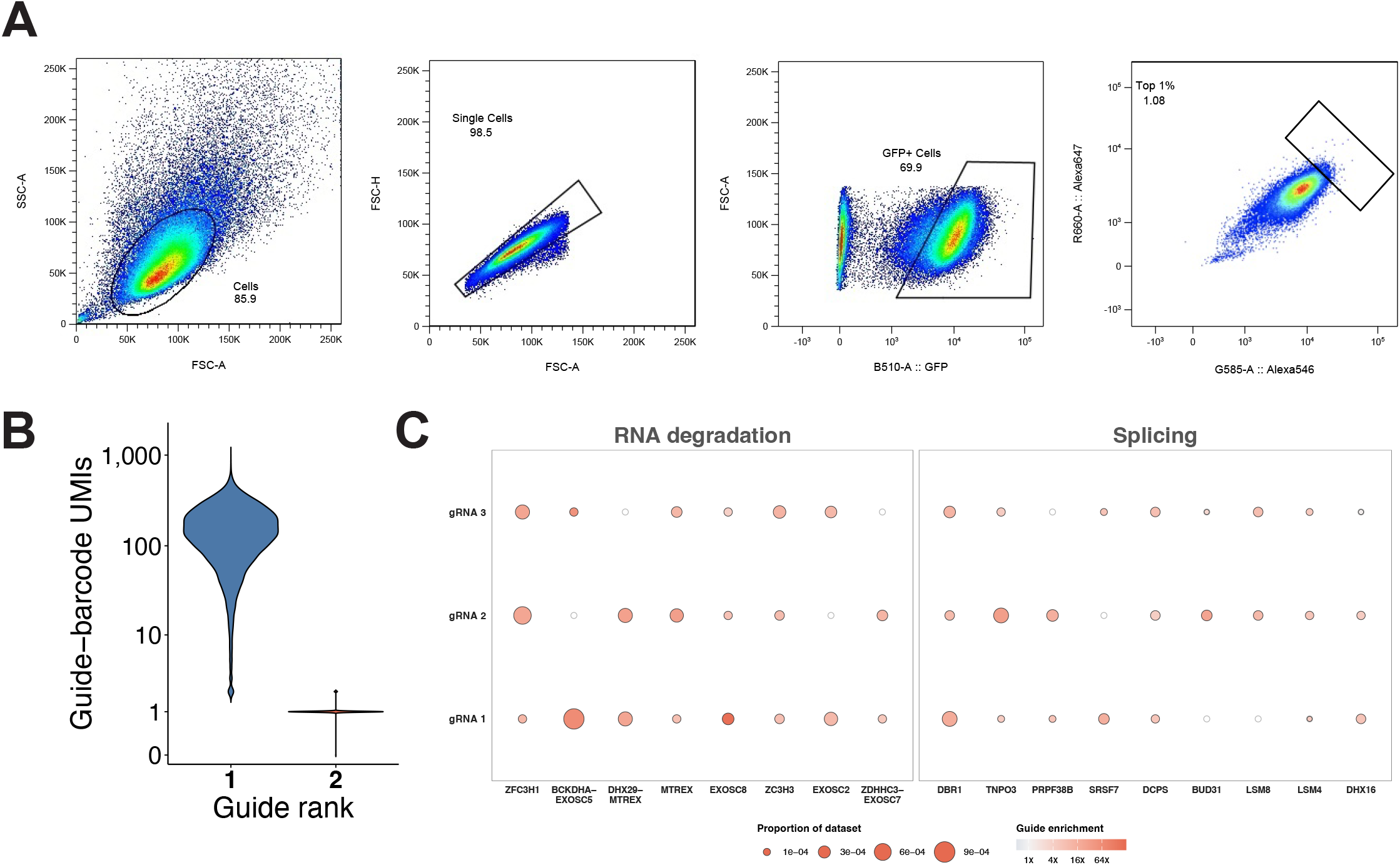
Quality control and guide-level reproducibility for the ncRNA VIPerturb-seq screen. **A)** Sequential FACS gating strategy used to isolate cells, singlets, GFP-positive cells, and the enriched population containing the top 1% of cells by combined *DANCR* and *SNHG3* fluorescence. **B)** Distributions of guide-barcode UMI counts for the highest-abundance (rank 1) and second-highest-abundance (rank 2) guides across the 21,644 gRNA singlets retained for analysis. A pseudocount of 0.5 was added only for visualization on the logarithmic axis. **C)** Guide-level enrichment for selected RNA-degradation and splicing hits, with genes shown along the x-axis and individual guides ordered by rank along the y-axis. Circle area represents the fraction of singlets assigned to each guide, and color represents enrichment relative to guide abundance in the plasmid library.

## SUPPLEMENTARY METHODS

### Cell culture

HEK293FT cells were acquired from Thermo Fisher (R70007). Monoclonal K562, HAP1, and HEK293 cell lines expressing KRAB-dCas9-MeCP2 (CRISPRi) ^51^ were derived as previously reported ^47^. K562 and HAP1 cell lines were maintained in Iscove’s Modified Dulbecco’s Medium (Thermo Scientific 31980030) supplemented with 10% FBS. HEK293 cell lines were maintained in High Glucose DMEM (Cytiva SH30243.01) supplemented with 10% FBS. All cells were grown at 37 °C and 5% CO_2_.

### Generation of GuEST-List barcode sequences

To generate a large pool of combinatorial barcodes to be detected by Flex probes, we designed matched libraries of LHS and RHS sequences. Barcode candidates were first generated by enumerating random sequences constrained by nucleotide patterns designed to balance GC content, prevent homopolymer runs, and match probe design constraints specified by 10x Genomics. Specifically, in IUPAC nucleotide code, the barcodes conformed to the following patterns:

LHS: NNNSWNNNSWNNNSWNNNSWNNNST

RHS: NNNWSNNNWSNNNWSNNNWSNNNWS

A minimum Hamming distance of 3 was enforced across the 10-base subsequences closest to the junction between the two probes (the 5^*′*^ end of the RHS barcode and the 3^*′*^ end of the LHS barcode), and an additional Hamming distance threshold of 8 was applied across the full length of each barcode. Sequences predicted to fold into secondary structures with Δ*G < −* 3 kcal*/*mol and with *T*_*m*_ *>* 65 °C using the Primer3 package were excluded ^22^. Remaining sequences were aligned to the NCBI RefSeq RNA database using BLASTn-short, and any barcode with fewer than five mismatches to a human or mouse RefSeq transcript, anti-sense RNA, or repetitive element was discarded. From this filtered set, a final pool of 238 LHS barcodes and 240 RHS barcodes was selected. Barcodes were then concatenated with constant regions required for probe synthesis, including capture sequences compatible with 10x Genomics protocols and amplification handles for library preparation and ordered as an oPool (Integrated DNA Technologies). Finally, each guide in the genome-wide library was assigned at random to a unique LHS-RHS barcode pair and one of three variable sgRNA scaffold sequences ^2,15^. These guidescaffold-barcode cassettes were concatenated with flanking sequences for downstream cloning into a CROPseq-style vector and ordered as an oligo pool from Twist Biosciences for cloning. The final library supports detection of up to 238 *×* 240 = 57,120 unique sequences.

### Cloning of GuEST-List

The vector backbone was prepared by digesting a modified CROPseq plasmid with Esp3I and BamHI restriction enzymes at 37 °C for 2 hours, to excise the entire sgRNA protospacer and constant region from the backbone. The cut plasmid components were purified with a silica column– based kit (Zymo D4005), and the backbone portion was isolated by performing a gel cut from a 1% agarose gel, using a commercially available gel purification kit (Zymo D4007), and performing a 2x SPRI bead clean-up. The insert library obtained from Twist (see “Generation of GuEST-List barcode sequences”) was amplified with 6 cycles of PCR according to the manufacturer’s instructions, using custom primers extending the region of homology with the modified CROPseq backbone. The product was purified with a 1.5x SPRI bead clean-up. The pooled insert and backbone were combined in a 5:1 ratio and incubated with Gibson Assembly Mix (New England Biolabs E2611L) as indicated by the manufacturer at 50 °C for one hour. The resulting plasmid library was then purified with isopropanol precipitation, dissolved in TE buffer, and used to transform Endura electrocompetent cells (Lucigen 60242-1). Transformed bacteria were cultured on LB-agar plates supplemented with ampicillin, and allowed to grow overnight at 37 °C. The final plasmid library was isolated by maxiprep (IBI Scientific IB47120) and validated by whole-plasmid sequencing (Azenta).

### Individual sgRNA cloning

For validation of individual hits, sgRNA constructs were prepared as previously described ^52^. Briefly, two oligonucleotides corresponding to the protospacer of each guide with overhangs were ordered from IDT as specified in Supplementary Table 8 and diluted to 100 *µ*M. For each protospacer, 1 *µ*L of each oligo strand was incubated together with T4 PNK (New England Biolabs M0201S) in T4 ligation buffer (New England Biolabs B0202S) and incubated at 37 °C for 30 minutes, then 95 °C for 5 minutes followed by a 5 °C min^*−*1^ ramp down to 25 °C. A modified CROPseq plasmid was digested with Esp3I restriction enzyme at 37 °C for 2 hours to excise the sgRNA protospacer stuffer sequence, incubated for an additional 45 minutes with FastAP phosphatase (Thermo Fisher EF0654) and purified via gel cleanup as previously described (see Cloning of GuEST-List). The annealed oligo insert and backbone were ligated together at a 10:1 ratio with T4 ligase (New England Biolabs M0202S) in T4 ligation buffer overnight at 4 °C, before being transformed into Stbl3 cells with heat shock at 42 °C for 45 seconds and cultured overnight on LB-Agar plates supplemented with ampicillin. Individual colonies were picked and grown overnight at 37 °C with shaking in LB-Amp media. Plasmid DNA for each construct was isolated via miniprep (Qiagen 27106).

### Lentivirus generation

Virus was prepared for transduction of human cell lines with the GuEST-List construct using standard second-generation lentiviral production methods. Briefly, HEK293FT cells were seeded with High Glucose DMEM (Cytiva SH30243.01) + 10% FBS (D10) in 10 cm dishes to achieve confluency between 85-90% at the time of transfection. Packaging and envelope plasmids psPAX2 (6 *µ*g; Addgene #12260) and pMD2.G (3 *µ*g; Addgene #12259) and GuEST-List plasmid (9 *µ*g) were co-transfected into the HEK293FT cells using Lipofectamine 3000 (Thermo Scientific L3000008). After 6 hours of incubation, the transfection medium was refreshed with D10 supplemented with 1% BSA (Thermo Scientific J6410018). The viral supernatant was harvested at 24 hours and 72 hours after transfection, passed through a 0.45 *µ*m filter, precipitated overnight at 4 °C (Alstem Cell Advancements VC100), concentrated in 100 *µ*L of PBS, and stored at *−*80 °C.

### Lentiviral transduction and cellular selection

Lentivirus was titrated in target cell lines over a range of 1 *µ*L - 10 *µ*L per 3 million cells to determine the optimal dosage for achieving an MOI within the range 0.3-0.5. For VIPerturb-seq experiments, cells were transduced at 50x library coverage in 12-well plates using aliquots containing 3 million cells in 2 mL. Lentivirus was allowed to incubate with cells for 24 hours before the cells were pooled and transferred to a fresh vessel with appropriate medium supplemented with 1 *µ*g mL^*−*1^ puromycin for HEK293 and HAP1 cells, and 2 *µ*g mL^*−*1^ puromycin for K562 cells. For the duration of chemical selection, the medium was replaced with fresh puromycin-supplemented medium every 24 hours.

### Antibody conjugation

Antibody conjugation was performed as previously described ^53^. Briefly, single-stranded oligonucleotides with a 5^*′*^ amino modifier C12 were ordered from IDT at a 100 nmol synthesis scale (sequences are available in Supplementary Table 3). A TCO-PEG4 linker was attached to each oligo through incubation with 0.5 mM TCO-PEG4-NHS reagent (Click Chemistry Tools A137-25) and column purified with Micro Bio-Spin 6 Columns (Bio-Rad 732-6221). Purified antibody was labeled with mTz-PEG4 (Click Chemistry Tools 1069-10) and conjugated to the TCO-PEG4 labeled oligo at a target concentration of 15 pmol oligo per 1 *µ*g of antibody. For hashing antibodies, 10 *µ*g of each antibody was passed through a 50 kDa MWCO filter (EMD Millipore UFC505024) 3 times. For intracellular staining panels, 3 *µ*g of each antibody was pooled together and treated with 40% ~4.32 M saturated ammonium sulfate, then passed through a 50 kDa MWCO filter 3 times.

### Multimodal cell mixing pilot experiment

#### Fixation and permeabilization

K562-, HAP1-, and HEK293-CRISPRi cell lines were transduced with the pilot library virus and subjected to puromycin selection as described above. At 10 days of selection, cells were harvested and FlexPlex was performed as previously described ^19^ with modifications. Briefly, 1 million cells per cell line were washed 1x with 1 mL PBS, then resuspended in 450 *µ*L of PBS and passed through a 40 *µ*m filter. 30 *µ*L of 16% formaldehyde solution (Sigma F8775-25ML) was added for a final concentration of 1% formaldehyde and allowed to fix at room temperature for 10 minutes. The fixation was quenched by the addition of glycine to a final concentration of 125 mM, and washed 2x with 1 mL of PBS. Cells were then resuspended in 100 *µ*L of cold lysis buffer (10 mM Tris-HCl, 10 mM NaCl, 3.33 mM MgCl_2_, 0.1% NP-40 (Thermo 28324), 1% BSA, 0.1 mM DTT (Invitrogen y00147), 200 U NxGen RNase Inhibitor (Lucigen 30281-1)) and allowed to incubate on ice for 5 minutes. Once lysis was complete, 1 mL wash buffer #1 (10 mM TrisHCl, 10 mM NaCl, 3.33 mM MgCl_2_, 1% BSA, 0.1 mM DTT) was added to the samples, and the cells were pelleted with a 5-minute spin at 500 rcf.

#### Hashing and intracellular antibody staining

Cells were then resuspended in 100 *µ*L of blocking buffer (1x PBS supplemented with 3% BSA, 100 *µ*g random blocking oligo ([Nx30]/3ddC/), 100 U NxGen RNase inhibitor, and 0.1 mM DTT). 1 *µ*g hashing antibody prepared as described above was added to each sample, and cells were allowed to incubate for 30 minutes at room temperature with rotation. Cells were then washed twice in 1x PBS supplemented with 3% BSA and 0.1% Tween-20, pooled together, and washed once more.

During wash steps, the intracellular antibody pool prepared as described above was incubated with 8 *µ*g of singlestranded DNA binding protein (Promega M3011) per 1 *µ*g of antibody in a solution of 1x NEB buffer 4 (NEB B7004S) for 30 minutes at 37 °C. After completing the incubation, BSA and PBS were added to a final concentration of 3% BSA in 1x PBS. Cells were resuspended directly in the antibody solution, and allowed to incubate for 1 hour at room temperature with rotation.

#### 10x Genomics Flex v1 multimodal workflow

To preserve antibody binding, cells underwent a secondary fixation step for one hour at room temperature following the 10x Genomics recommended protocol (CG000782). After the additional fixation, cells were taken directly as input into the 10x Genomics Flex v1 protocol (CG000786), with the following modifications: While preparing the probe hybridization reaction (step 1.1.g), 2.5 *µ*L of a 40 nM working stock of the custom guide-barcode pilot probe set was spiked in, according to 10x Genomics recommendations for custom probes (CG000621). When preparing the cell dilution for GEM formation (step 3.1.b), 40,000 cells were loaded per lane, superloading above official kit guidelines ^54^. When setting up the preamplification PCR reaction (step 4.2.a), 1 *µ*L of 0.2 *µ*M ADT Additive Primer, GDO Additive Primer, and HTO Additive primer were each added to the reaction (full sequences for all custom primers used in the Flex workflows are available in Supplementary Table 4).

#### Library preparation

After completing step 4.3.o of the 10x Genomics Flex v1 protocol, 100 *µ*L of the cleaned-up preamplification material was divided among separate library preparations for each modality as follows:

For the gene expression library, 20 *µ*L of preamplification was combined with 50 *µ*L of Amp Mix C (10x Genomics PN 2001311), 25 *µ*L nuclease-free water, 2.5 *µ*L of 10 *µ*M Universal Sample Index Forward Primer, and 2.5 *µ*L of 10 *µ*M GEX Sample Index Reverse Primer. The PCR reaction was performed under the following conditions: 98 °C for 45 s, (98 °C for 20 s, 54 °C for 30 s, 72 °C for 20 s) x 6 cycles, 72 °C for 1 min. The final library was purified with a 1.0x SPRI clean-up and eluted into 41 *µ*L Buffer EB (Qiagen 19086).

For the antibody-oligo libraries, 20 *µ*L of preamplification was combined with 50 *µ*L of 2x KAPA HiFi HotStart ReadyMix (Roche 07958935001), 25 *µ*L nuclease-free water, 2.5 *µ*L of 10 *µ*M Universal Sample Index Forward Primer, and 2.5 *µ*L of 10 *µ*M HTO Sample Index Reverse Primer (for hashing antibodies) or 10 *µ*M ADT Sample Index Reverse Primer (for intracellular protein library). The PCR reaction was performed under the following conditions: 95 °C for 3 min, (95 °C for 20 s, 60 °C for 30 s, 72 °C for 20 s) x 14 cycles, 72 °C for 5 min. The final library was purified with a 1.6x SPRI clean-up and eluted into 21 *µ*L Buffer EB.

For the guide-barcode libraries, an enrichment PCR was first performed by combining 20 *µ*L of preamplification material with 50 *µ*L of Amp Mix C, 25 *µ*L nuclease-free water, 2.5 *µ*L of 10 *µ*M Universal Sample Index Forward Primer, and 2.5 *µ*L of 10 *µ*M GDO Enrichment Reverse Primer. The PCR reaction was performed under the following conditions: 98 °C for 45 s, (98 °C for 20 s, 54 °C for 30 s, 72 °C for 20 s) x 9 cycles, 72 °C for 1 min. The enriched library was purified with a 1.8x SPRI clean-up and eluted into 30 *µ*L Buffer EB. This enriched library was combined with 50 *µ*L of 2x KAPA HiFi HotStart ReadyMix, 25 *µ*L nuclease-free water, 2.5 *µ*L of 10 *µ*M Universal Sample Index Forward Primer, and 2.5 *µ*L of 10 *µ*M GDO Sample Index Reverse Primer to set up a final, sample-indexing PCR reaction, performed under the following conditions: 95 °C for 3 min, (95 °C for 20 s, 60 °C for 30 s, 72 °C for 20 s) x 8 cycles, 72 °C for 5 min. The final library was purified with a 1.6x SPRI clean-up and eluted into 21 *µ*L Buffer EB.

### Recombination benchmarking experiment

Nine sgRNAs inducing strong perturbations and one nontargeting sgRNA were individually cloned following the protocol specified above (see “Individual sgRNA cloning”). Individual colonies were picked and grown overnight at 37 °C with shaking in LB-Amp media. Plasmid DNA was isolated from each culture via miniprep (Qiagen 27106); the guide– barcode cassette was verified by Sanger sequencing from the U6 promoter (Azenta); and the constructs were manually pooled. Virus was prepared from the resulting plasmid pool as previously described (see “Lentivirus generation”). K562-CRISPRi cells were transduced with this benchmarking virus and subjected to puromycin selection as described above for 8 days. After selection, cells were harvested and 300,000 cells were input into a single hybridization reaction using the 10x Flex v2 workflow as described above (see “Genome-wide single-cell screen with GuEST-List and Flex v2”) with minor modifications. When preparing probe hybridization reactions (step 1.1.b), 2.75 *µ*L of 40 nM custom probes against the guide protospacers on the Nextera R2 handle were spiked into the whole transcriptome hybridization reaction along with 2.75 *µ*L of 40 nM custom guide-barcode probes according to the 10x Genomics recommendations. Libraries for protospacer probes were prepared following the modified antibody-oligo library protocol described above, as both libraries were on the Nextera R2 sequencing handle, and all other libraries were prepared for sequencing as described above (see “Multimodal cell mixing pilot experiment”).

Separate cell-by-guide UMI count matrices were generated for the protospacer and guide-barcode libraries. Guide identities were assigned independently for each assay using `fishash`^55^ with an adjusted *P*-value threshold of 0.05. To estimate the recombination rate, we retained cells classified as singlets by both assays and required at least 80% of protospacer UMIs to correspond to the dominant protospacer assignment, yielding 7,562 high-confidence cells. For fingerprint analysis, gene-expression counts were normalized using SCTransform, and features were selected independently for protospacer- and barcode-based assignments using `ChooseFeatures` with the non-targeting guide as the control. The intersection of the two feature sets was used to learn separate perturbation dictionaries with `LearnDictionary`. Concordance was assessed using Pearson correlations between the matched protospacer- and barcode-derived fingerprint vectors, and the median correlation across perturbations was 0.97.

### Genome-wide single-cell screen with GuEST-List and Flex v2

K562-CRISPRi cells were transduced with CRISPRi GuEST-List GFP v1 and subjected to puromycin selection as described above. After 6 days of selection, 20 million cells were harvested, washed 1x in PBS and resuspended in MACS buffer. Fluorescence-activated cell sorting (FACS) was performed on a Sony SH800 instrument with a 100 *µ*m chip to isolate 10 million GFP+ singlets. These cells were input directly into the 10x Genomics recommended fixation and permeabilization workflow (CG000782). From there, cells were split into 24 aliquots of 300,000 each and directly input into the 10x Genomics GEM-X Flex v2 workflow for Multiplexed samples (CG000834) with the following modifications: When preparing probe hybridization reactions (step 1.1.b), 66 *µ*L of 40 nM custom probe working stock was spiked into the 24-reaction probe hybridization mix, as per the official 10x guidelines for the use of custom probes with Flex v2 (CG000839). When setting up the preamplification reactions (step 5.2.a), 1 *µ*L of 10 *µ*M custom GDO Additive v2 Primer was spiked in. To prepare the final indexed libraries, 20 *µ*L of preamplified material from step 5.3.p was split across four separate PCR reactions each containing 50 *µ*L Amp Mix C, 30 *µ*L of nuclease-free water, 5 *µ*L of 10 *µ*M Universal Sample Index Forward Primer, and 5 *µ*L of 10 *µ*M custom GEX Sample Index Reverse Primer (for the whole transcriptome library) or 5 *µ*L of 10 *µ*M custom GDO Sample Index v2 Reverse Primer (for the guide-barcode library). PCR was performed under the following conditions: 98 °C for 45 s, (98 °C for 20 s, 54 °C for 30 s, 72 °C for 20 s) x 5 cycles (gene expression) or 13 cycles (guide-barcode), 72 °C for 1 min. Final libraries were purified with a 1.0x SPRI clean-up and eluted in 40 *µ*L Buffer EB.

### VIPerturb-seq for regulators of vimentin

#### Fixation and permeabilization

K562-CRISPRi cells were transduced with CRISPRi GuEST-List GFP v1 and subjected to puromycin selection as described above. After 10 days of selection, 40 million cells were harvested, washed 1x in PBS-T and filtered through a 40 *µ*m strainer. Cells were then resuspended in a solution of 4% paraformaldehyde in PBS and allowed to fix for 1 hour at room temperature. After one hour, cells were pelleted and washed twice in PBS-T. Cells were then resuspended in ice-cold 70% ethanol and allowed to permeabilize overnight at 4 °C. After incubation, cells were spun down at 850 rcf and washed twice in PBS-T.

#### Intracellular staining and phenotypic enrichment

Cells were next resuspended in blocking buffer and allowed to incubate for 30 minutes at room temperature with rotation. Cells were then washed twice with PBS-T, resuspended in staining buffer supplemented with 1 *µ*L of vimentin-AF647 antibody (BioLegend 677807) in 100 *µ*L of buffer per million cells and allowed to incubate for one hour at room temperature with rotation. FACS was performed on a Sony SH800 instrument with a 100 *µ*m chip. Cells were gated to isolate GFP+ singlets in the top 3% of the distribution for vimentin fluorescence.

#### 10x Genomics Flex v1 Workflow

100,000 cells were recovered after sorting and used directly as input into the 10x Genomics Flex v1 workflow (CG000786), with the modifications described above for the multimodal cell-line mixing pilot experiment, omitting the inclusion of primers and additional library preparation steps for antibody-oligo modalities that were not included in this experiment.

### VIPerturb-seq for regulators of ncRNA abundance

#### Custom ncRNA probes

Custom probes were designed against 30 ncRNAs not included in the default human whole transcriptome probe set, according to recommendations from 10x Genomics. Full sequences for all custom ncRNA probes can be found in Supplementary Table 7.

#### HCR™ FlowFISH

K562-CRISPRi cells were transduced with CRISPRi GuEST-List GFP v1 and subjected to puromycin selection as described above for 6 days. After selection, 75 million cells were harvested, enriched for live cells (Miltenyi 130-090-101) and subjected to the 4% paraformaldehyde fixation and 70% ethanol permeabilization protocol described above (see “VIPerturb-seq for regulators of vimentin”). After permeabilization, HCR™ FlowFISH was performed on the cells with probes against *DANCR* and *SNHG3* ncRNA transcripts according to the previously described PERFF-seq protocol ^12^. In brief, 40 million cells were washed twice with 40 mL of PBS-T and subsequently incubated with 4 mL of HCR™ HiFi Probe Hybridization Buffer at 37 °C and 300 rpm shaking for 30 minutes. 80 *µ*L of each probe solution targeting *DANCR* and *SNHG3* was diluted in 4 mL HCR™ HiFi Probe Hybridization Buffer and added to the cells. Cells were incubated for an additional 3 hours at 37 °C with shaking. After hybridization, cells were washed with 20 mL HCR™ HiFi Probe Wash Buffer 3 times. Cells were next incubated in 4 mL Gold Amplifier Buffer for 30 minutes at room temperature. 160 *µ*L of each hairpin was aliquoted, heated to 95 °C for 90 seconds, and then allowed to cool to room temperature in a dark drawer for 30 minutes. The cooled hairpins were diluted in 100 *µ*L HCR™ Gold Amplifier Buffer/million cells and added to the sample. Amplification was allowed to proceed overnight at room temperature in a tube rotator protected from light. Cells were subsequently washed seven times with HCR™ Gold Amplifier Wash Buffer before resuspending in PBS-T supplemented with 1% BSA and filtering through a 40 *µ*m mesh.

FACS was performed on a FACSSymphony S6 instrument. Cells were gated to isolate GFP+ singlets in the top 1% of the distribution for *DANCR* and *SNHG3* fluorescence. 100,000 cells were recovered after sorting and stored at 4 °C according to guidelines from 10x Genomics. A DNase digestion step was performed immediately after sorting to strip bound FlowFISH probes and amplified hairpins, which can interfere with the 10x Flex reactions as previously described ^12^.

#### 10x Genomics Flex v2 workflow

After probe stripping, cells were input into the 10x Genomics Flex v2 workflow with the modifications described above for the genome-wide single-cell screen with Flex v2. For this experiment, we included custom probes against a small set of ncRNAs (Supplementary Table 7), which were mixed in with the custom GDO probes at the same concentration and amplified alongside the whole transcriptome.

### FlowFISH of single perturbations

Lentivirus was prepared in an arrayed fashion from single-guide libraries cloned for each potential hit (see “Individual sgRNA cloning”). HCR™ FlowFISH was performed as previously described, with an input of 1 million cells per condition on a Sony SH800 instrument. The data were analyzed in FlowJo v10.10.0 and R v4.5.2.

Statistical significance was assessed using a linear model fitted to the percentage of RNA-positive cells across all conditions. The model included RNA target, perturbation, their interaction, and experimental replicate as fixed effects, thereby estimating perturbation- and RNA-specific means while pooling residual variance across the complete experiment. Within each RNA target, two-sided model contrasts compared each perturbation with the non-targeting control.

### Sequencing, preprocessing, and QC

For Flex v1 experiments, libraries were sequenced using a NextSeq 550 instrument at a depth of approximately 10,000 gene expression reads per cell, 1,000 guide-barcode reads per cell, and (for the multimodal cell-line mixing pilot only) 1,000 antibody-oligo reads per cell. For genome-wide experiments, libraries were sequenced using an Ultima UG100 instrument at a depth of approximately 25,000 gene expression reads per cell and 700 guide-barcode reads per cell. Raw unaligned CRAM files were converted to FASTQs using SAMtools ^56^. Adapter trimming and read formatting were performed with cutadapt ^57^ to generate FASTQs compatible with alignment and quantification using Cell Ranger 9.0. For the ncRNA VIPerturb-seq screen, libraries were sequenced on an Element AVITI instrument.

Analysis of all single-cell datasets was performed using Seurat v5.3.1 ^58^. For all single-cell experiments, Seurat objects were filtered for cells passing a UMI threshold of 1000. To quantify guide-barcode expression, we constructed a cell-by-guide UMI count matrix directly from the guide-barcode FASTQ files. Briefly, we iterated through all reads in the guide-barcode FASTQ and extracted the associated cell barcode, guide-barcode sequence, and UMI for each read. Reads were first filtered to retain only those whose cell barcode matched a barcode identified as a valid cell in the corresponding Cell Ranger gene expression output, allowing up to one base mismatch to account for sequencing errors. For retained reads, the guide-barcode sequence was then matched to the reference guide-barcode list, again permitting up to one base mismatch. UMIs were tracked on a per-cell, per-guide basis, and only previously unseen UMIs were counted. This procedure yielded a UMI count matrix representing guide-barcode abundance for each cell. We loaded this matrix into Seurat, and used MULTISeqDemux ^59^ to demultiplex data and assign gRNA identity to individual cells, retaining cells that were assigned as gRNA singlets.

### Analyzing perturbations with RNA fingerprinting

We analyzed Perturb-seq datasets using our previously developed RNA fingerprinting framework ^20^. This framework performs two complementary tasks. First, for each perturbation profiled in the VIPerturb-seq dataset, it learns a transcriptional fingerprint: a genome-wide vector that quantifies the effect of the perturbation on the expression of every measured gene. We have previously shown that this vector is more informative than traditional differential expression statistics, as it accounts for heterogeneity between cells driven by either differences in baseline state, or differences in perturbation efficiency ^20^.

Second, the framework enables systematic matching of learned fingerprints to a reference Perturb-seq dictionary, allowing perturbations profiled in VIPerturb-seq to be compared against previously characterized genetic perturbations. Fingerprint matching was visualized using Long Island City plots, which summarize similarity relationships between perturbations, and with ‘matches’ denoted in red. We applied this approach to compare VIPerturb-seq perturbations to reference fingerprints derived from both the FlexPlex dataset ^19^ and the Genome-wide Perturb-seq dataset ^5^.

We used the `LearnDictionary, FingerprintCells`, and the `LongIslandCityPlot` functions from the RNA fingerprinting package with default parameters for these analyses. For the pilot experiments, we used RNA fingerprinting to confirm that perturbations measured by VIPerturb-seq correctly mapped to perturbations measured by previously collected reference Flex Perturb-seq datasets where custom probes directly capture the gRNA protospacers ^19^.

For the genome-wide experiment with Flex v2, we estimated RNA fingerprints for each of the perturbations in the GuEST-List library. For all perturbations, we examined the fingerprint vector’s value for the target gene (i.e., asking how CRISPRi perturbation of SMG5 affects the expression of SMG5), and observed a negative value in 87.7% of cases. The remaining cases may represent inefficient knockdown across all gRNAs, or insufficient data to infer a perturbation-driven reduction in a single gene.

We also performed unsupervised clustering analysis of perturbations to reveal groups of functionally related regulators. For this analysis, we first performed principal components analysis on the fingerprint matrix, projecting each perturbation into a 50-dimensional space. We calculated a correlation matrix of all fingerprints in this space, and removed all perturbations that did not exhibit correlation with another perturbation of *>* 0.6. We further removed perturbations that did not exhibit correlation with another perturbation in the original space (prior to PCA) of *>* 0.2. For the remaining 667 perturbations, we obtained a clustered correlation matrix using the ComplexHeatmap package, and manually annotated 21 functionally enriched correlated modules.

For the single-cell heatmaps in Figure 3I, we visualized expression profiles for 10 up-regulated genes for perturbations previously known to represent 10 regulatory processes. We selected the top up-regulated genes based on the ten largest values in the fingerprint matrix for each perturbation. We used Mixscape ^7^ to remove ‘escaping’ cells that did not exhibit transcriptomic evidence of receiving a perturbation despite receiving a gRNA. We did not run Mixscape for any other analyses in this manuscript.

To determine perturbation enrichment in the VIM regulator experiment, we first estimated the background gRNA abundance in the GuEST-List plasmid library. To do this, we sequenced the pooled plasmid library used for the VIPerturb-seq screen prior to cell transduction. For each perturbation, the expected background frequency was calculated as the fraction of total plasmid reads corresponding to each gene. The abundance in the phenotypically enriched VIPerturb-seq dataset for each perturbation was calculated as the fraction of the total number of sequenced cells assigned to a gene perturbation, divided by the total number of cells in the library. The normalized cell abundance shown in Figure 4B is the ratio of these two values.

We also assessed the empirical enrichment by comparing the observed cell counts in the genome-wide screens performed with and without FACS enrichment (Figure 4C). We calculated the proportion of cells assigned to each perturbation in each experiment, averaging across gRNAs (up to *n* = 3 for targeting guides, *n* = 500 for non-targeting controls). The ratio between these two numbers represents the degree of enrichment.

### Computational analysis of the ncRNA screen

To calculate enrichment in the ncRNA-high population, we counted the number of selected cells assigned to each sgRNA and compared this value with the abundance of that sgRNA in the plasmid library (adding a pseudocount of one for each sgRNA). Gene-level enrichment was defined as the mean enrichment across sgRNAs targeting the same gene.

To obtain a molecular phenotype score, we used SCTransform v2 ^60,61^ to estimate Pearson residuals for *SNHG3* and *DANCR*. For each targeted gene, we summed the *SNHG3* Pearson residuals across all selected cells carrying an sgRNA targeting that gene and separately summed the corresponding *DANCR* residuals. These two sums were then added to define the combined molecular phenotype score. Genes represented by fewer than three selected cells were excluded. Enrichment and molecular phenotype values were independently converted to percentile ranks, and the mean of the two percentile ranks was used as a joint-prioritization score. Genes were ranked in descending order of this score for Figure 5E and subsequent analyses.

For gene ontology analysis, we selected the 50 genes with the highest joint-prioritization scores. Unique genes in the resulting list were submitted to Enrichr using the GO Biological Process 2025 library. Enrichment significance was assessed using Benjamini–Hochberg-adjusted *P* values returned by Enrichr.

To reduce redundancy among enriched terms, terms were ordered by adjusted *P* value and then nominal *P* value. A term was considered redundant when at least 50% of the smaller of its overlapping-gene set and that of a previously retained, more significant term was shared. Figure 5F displays the two retained biological processes.

For Figure 5H, log-normalized expression values for *SNHG3* and *DANCR* were aggregated separately for each sgRNA. Each point therefore represents one sgRNA with at least one cell in the selected population. Guide-level values for splicing and RNA-degradation perturbations were compared separately with non-targeting guides using two-sided Wilcoxon rank-sum tests. Boxplots summarize the guide-level distributions, and dashed horizontal lines denote the corresponding non-targeting medians.

